# Attributing the variability in direction and magnitude of local-scale marine biodiversity change to human activities

**DOI:** 10.1101/162362

**Authors:** Jillian C Dunic, Robin Elahi, Marc J. S. Hensel, Patrick J. Kearns, Mary I. O’Connor, Daniel Acuña, Aaron Honig, Alexa R. Wilson, Jarrett E. K. Byrnes

## Abstract

In recent decades, environmental drivers of community change have been associated with changes in biodiversity from local to global scales. Here we evaluate the role of anthropogenic drivers in marine ecosystems as drivers of change in local species richness with a meta-analysis of a novel dataset of temporal change in species richness. We paired biodiversity data from 144 sites with large-scale drivers derived from geospatial databases: human cumulative impact scores, sea surface temperature change, nutrient loading, and invasion potential. Three specific drivers (nutrient inputs, rate of linear temperature change, and non-native species invasion potential) explained patterns in local marine species richness change. We show that these drivers have opposing effects on biodiversity trends. In some cases, variability in drivers can create contrasting directions of change yielding observations of no net change when localities are pooled in an attempt to find a global average. Further, long-term studies reveal different effects of drivers that are not observed in short-term studies. These findings begin to explain high variability observed in species diversity trends at local scales. Formally attributing local species diversity change to human drivers is essential to understanding global patterns of local species diversity change and their consequences.

## INTRODUCTION

Human impacts such as habitat destruction, pollution, and climate change have reduced global species diversity (Vié et al. 2008, Newbold et al. 2015). At the same time, local temporal trends in diversity are variable; syntheses of marine and terrestrial diversity time-series have reported decreases, increases (Elahi et al. 2015, Schipper et al. 2016), and no net change in local scale (e.g., <20 km^2^) species richness (Vellend et al. 2013, Dornelas et al. 2014) over the last century. These syntheses include such wide variation as observations ranging from 8% species loss/yr to 35% species gains/yr in plant communities (Vellend et al. 2013) and 5% species loss/yr to 6% species gains /yr in coastal marine communities (Elahi et al. 2015). This variation in temporal trends suggests that understanding global trends in biodiversity requires understanding local community change and the different interactions with human drivers in different places and at different scales (McGill et al. 2014, Elahi et al. 2015).

This attribution of human impacts to variability in trends is crucial to understanding future biodiversity change. We know human-mediated disturbances contribute to local species loss and gains across terrestrial (Murphy and Romanuk 2013, Newbold et al. 2015), aquatic (Lake et al. 2000), and marine environments (Dulvy et al. 2003, Sala and Knowlton 2006). Habitat change, overexploitation, and pollution negatively affect species at a local scale (Jackson et al. 2001, Jones et al. 2004, Lotze et al. 2006, Johnston and Roberts 2009). Further, cumulative stressors and their interactions can amplify negative community responses to human impacts [15,16]. In contrast, local diversity can increase due to human impacts such as invasions and climate change (Sax et al. 2002, Menendez et al. 2006). If multiple opposing drivers interact or co-occur, they could produce observations of no net change in local scale species richness. Elahi et al. (2015) and Vellend et al. (2013) attributed some variation in diversity trends to localized human activities as reported in the original studies. Space-for-time substitutions have been used to search for global patterns of local-scale diversity change in response to human drivers (Murphy and Romanuk 2013, Newbold et al. 2015). Still, no assessment of diversity temporal trends has examined the potential role of human drivers as a predictor of variation in trends in local diversity over time.

Here we attempt to attribute the effects of human impacts on patterns of local-scale species richness change in coastal marine communities. Coastal communities worldwide are subjected to a range of different human activities that reduce the abundance of many marine species (Halpern et al. 2008). We consider the possibility that cumulative human impacts reflect additive stresses on marine species and thus have a negative effect on local species richness (Crain et al. 2008). We then quantify how a combination of the human impacts—nutrient addition, shipping traffic (as a proxy for invasion potential), and rate of temperature change—affect species richness change in marine communities at a local/site level. These impacts could have positive or negative impacts on diversity. Nutrient addition can reduce local richness by degrading habitat (Gough et al. 2000, Conley et al. 2007) but can also increase productivity, and consequently increase total abundance and diversity. Shipping traffic facilitates species invasions and can lead to gains or offset local losses by introducing new species (Lockwood et al. 2005). Increased temperatures could lead to species range expansions yielding local gains of warm water species (Barry et al. 1995, Sorte et al. 2010, Burrows et al. 2014, Batt et al. 2017) if species dispersal rates can track changes in temperature (Pinsky et al. 2013). We expect that the observed effect of drivers on rate of change in diversity may differ in magnitude or sign depending on study duration reflecting differing effects of individual drivers over short versus long periods of time (Gonzalez et al. 2016). To test these predictions, we collated studies that have measured species richness from sites across marine biomes and leverage the available variation in global drivers to put our analysis in an ecological context. We find that variation in local-scale biodiversity change is related to the influence of human impacts such as climate change, invasions, and eutrophication.

## METHODS

### Study Selection

We systematically searched the literature using Web of Science and the Aquatic Commons database, which included grey literature publications. Additional grey literature publications were extracted from Elahi et al. (2015), which was published after our initial search. We searched for studies that had resampled marine species richness or diversity at a minimum of two time points equal to or greater than one year apart. Our literature search terms were adapted from Vellend et al. (2013) with keywords to target marine habitats while excluding freshwater or terrestrial habitats: e.g., ‘marine’ OR ‘ocean’ NOT (‘freshwater’ OR ‘terrestrial’), and combining these with keywords about biodiversity and resampling: (‘biodiv* OR ‘divers*’ OR ‘richness’) AND ‘resamp*’ (see full search string in Appendix S2). We initially entered the search terms into the Web of Science and Aquatic Commons databases on February 19, 2014. This search returned 4803 references, which we filtered down to 745 papers after reviewing titles, abstracts, and full text where necessary, to identify studies that met the following criteria: sampled marine taxa, reported biodiversity, and resampled sites with at least one year between initial and final sampling points (Appendix S1: Fig. S1). From the remaining 745 papers, we excluded studies if sampling methods were inconsistent between time points, if rare species were not included, or if authors identified events occurring before or during the study (e.g., Marine Protected Area implemented, resource extraction, construction) and had *a priori* expectations (see Appendix S2). After study selection we had a total of 144 time series from 115 sites and 34 studies globally (Appendix S1: Fig. S2). For taxonomic groups for which we had sufficient sample size, we tested the duration only and cumulative human impacts model, and found qualitatively similar results. However, the sample sizes in these analyses were small and we were unable to test individual drivers (see Appendix S3). Although we collected data for abundances (sites = 22) and Shannon diversity (time series = 48, sites = 40), the sample size was too small to evaluate the role of human impacts on this metric of diversity. Therefore, we discuss only results of local species richness change.

### Data Acquisition

We extracted 13 variables that described taxonomic group, sampling method, number of replicates, number of subsamples, plot size, and richness. In most studies, raw species abundance data were not provided and so data were extracted as summary statistics from figures using WebPlotDigitizer 3.10 (Rohatgi 2016) or manually extracted from data tables. Wherever possible, sampling errors from summary statistics were collected so that we could perform a variance-weighted meta-regression. When a site was sampled between a range of years (e.g., 1995 – 1996), the first year was recorded for consistency. When only a season or range of months was given, the average month of that season was recorded. We calculated the effect size as the log response ratio (LRR) of the proportion of species richness change between a final and initial time point (eqn. 1).

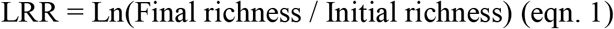

We chose to use the LRR instead of Hedges’s D, another commonly used metric of effect size, because log transformation of the response ratio normalizes the data and because we can use the following equation to convert the LRR into the percent change in species richness (eqn. 2).

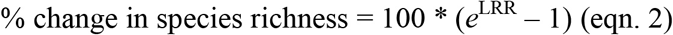

Zero values can be problematic when using log transformations, however, our data did not contain cases where species richness decreased to zero. We verified assumptions of normality of residuals for all models using visual inspection of standardized residuals and their quantiles. To check for potential publication bias in effect sizes, we visually inspected funnel plots. However, publication bias in our dataset was unlikely because many of our studies (46%) were not specifically testing for changes in biodiversity over time. For example, we used data from control sites from experiments where researchers were not examining change in ambient biodiversity over time *per se*. For a complete list of studies used in the analysis see Appendix S4.

### Driver data

To examine the effect of human impacts on the change in species richness over time, we used the cumulative human impacts (CHI) data created by Halpern et al. (2015). The CHI model summarizes data on a broad set of human impacts into a single score of cumulative human impact for every square kilometer of the world’s oceans. The scores are derived from a model that integrates global data for 19 different drivers including nutrient pollution, fishing, urban runoff, shipping traffic, and sea surface temperature anomalies (Halpern et al. 2015). The CHI model is a potential indicator of human impacts; however, this model incorporates the effects of multiple drivers that may have opposing effects on local species diversity. To also understand the effects of specific drivers on local-scale species richness change, we extracted data layers that had global coverage and that were expected to affect local richness in coastal areas. We used two data layers used in the CHI data: non-native species invasion potential (metric tonnes of cargo shipped to a port in 2011 was used as a proxy for invasion potential) and nutrient addition (metric tonnes of nitrogen and phosphorous fertilizer use as reported by the FAO from 2007 – 2010, was used as an indication of intensity of nutrient addition along coastal areas; See (Halpern et al. 2008, 2015 for details). We also calculated the decadal rate of linear temperature change (LTC), observed over the specific time span of each study, using the Met Office Hadley Centre Sea Surface Temperature data (Rayner et al. 2003). For each study, we collected the latitude and longitude of sampling points for all plots surveyed in a study. When study sites were composed of multiple subsamples, we included all the associated coordinates. Data from the spatial layers were then extracted from these coordinates. When a site was comprised of multiple coordinates, we computed the average impact value for each site.

### Statistical analysis

To examine whether marine species richness has changed at local scales and to test whether cumulative human impacts and specific drivers affect changes in local species richness, we performed three variance-weighted random effects meta-regressions using the package *metafor* (Viechtbauer 2010) in the statistical software R version 3.4.0 (R Core Team 2017). We included a random effect of study, as single studies could contain multiple sites. This approach allowed us to account for variation between studies due to factors such as differences in researcher methods, taxonomic groups, and sites. All code for analysis is available at https://github.com/jdunic/local-marine-meta.

To test the roles of specific drivers and to determine the average rate of change in local species richness we tested three models: the average change in local richness (eqn. 3), the effect of cumulative human impacts on local richness change (eqn. 4), and the effect of specific drivers on local richness change (eqn. 4). We used the model heterogeneity statistic Q_m_ to determine whether our models explained a significant amount of variability observed in the data. We first examined the average rate of change in species richness from our data set using the following model for site *i* from study *j*

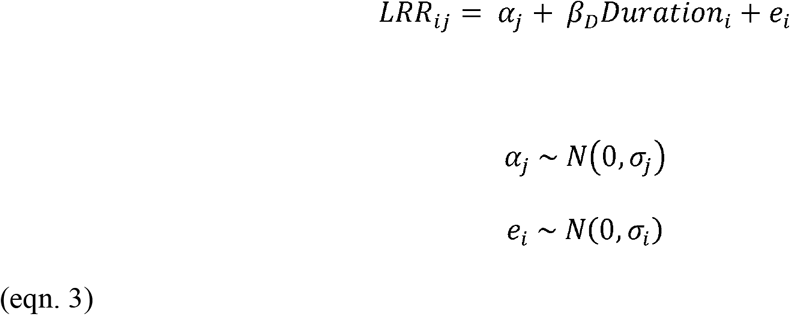

Where α_j_ is the between-study random effects (estimated by the model) and σ_i_ was the measured variance of a richness estimate at site *i*. We used study duration (years) as a predictor of the LRR to estimate a rate of change rather than use LRR / study duration as a response variable to increase the power of our analyses (Gonzalez et al. 2016). Thus, β_D_ describes the rate of change in the log response ratio of species richness (i.e., log change in richness per year). To ease interpretation, we can also convert β_D_ to a percent change in species richness per year (eqn. 2).

To evaluate the effects of different drivers, we used a general model for incorporating k drivers (eqn. 4).

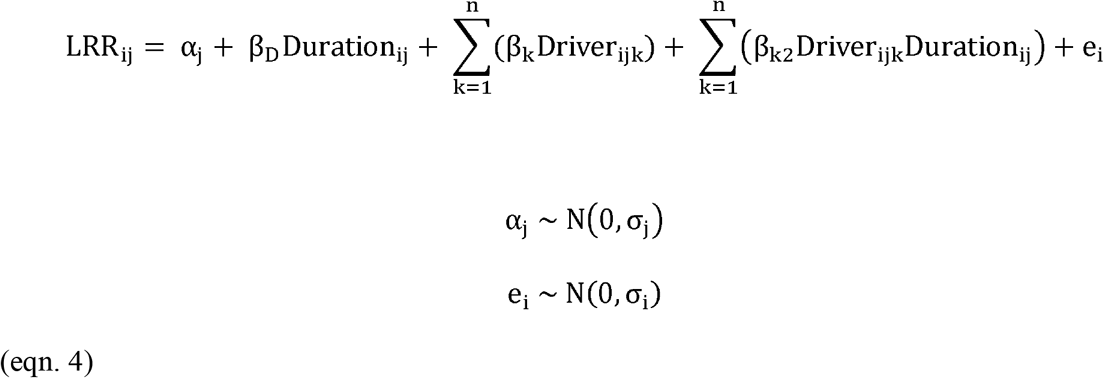

In this model formulation, the net log change in richness per year is now the sum of the baseline β_D_ for studies with zero human impacts and sum of all β_k2_ coefficients multiplied by their respective drivers. This allows us to account for differences observed in studies differing in duration by comparing the change in richness due to a driver irrespective of time (β_k_) – analogous to short-term changes due to a driver - versus how that driver affects the rate of log change in species richness (β_k2_). The latter can be seen as the effect of a driver that will be manifest over longer time-scales as the time independent component of an impact becomes less important. For example, a positive value for the coefficient β_k2_, would be interpreted as the driver slowing the rate of loss or increasing the rate of gain, depending on the sign of rate of change β_D_. Hence our model produced ‘long-term’ predictions of change due to a given driver by multiplying the effect out by, for example, 30 years. One limitation of this approach was that given our data, we only tested a linear effect of duration on how drivers affect rates of log richness change. Given the size of our data set this seemed a reasonable first approximation, although more model-based mechanistic studies could prove useful in the future. We used the Akaike Information Criterion corrected for small sample sizes (AICc) to compare models and determine whether inclusion of human impacts improved the predictive ability of the model relative to the model that included only study duration.

To determine whether any single study had a disproportionate effect on parameter estimates we systematically re-ran the meta-analysis excluding data from one study at a time to test the robustness of our analyses to outliers in the data (i.e., leave-one-out analyses, Appendix S4). We used variance weighting in our analysis because it increases the power to detect differences from zero by placing higher values on studies for which estimates are more precise (Lajeunesse 2013). Although using an unweighted or sample-size weighted analysis would increase the studies included in our analyses, the parameter estimates from these methods are unreliable (Appendix S1: Fig. S3). For completeness, however, we present these results along with their robustness to the exclusion of single studies using both methods (Appendix S4).

## RESULTS

### Local richness change

In general, inferences about the rate of change of species richness over time in our dataset depend on the inclusion of human impacts. In our model that did not account for human impacts, study duration influenced observed change in species richness (Q_m_ = 5.12, p = 0.024) and we found that on average, the log rate of change in species richness per year was 0.01 (95% CI = 0.002 – 0.021, p = 0.022, sites = 142, studies = 34), which corresponds to an average trend of species richness gains at a rate of 1.05% per year within our dataset (Fig. 1a, Appendix S1: Table S1). However, within our dataset, both models that included some form of human impacts performed better than the model that did not include human impacts (Table 1) and altered the mean estimated log ratio of species richness over time.

**Figure 1.**
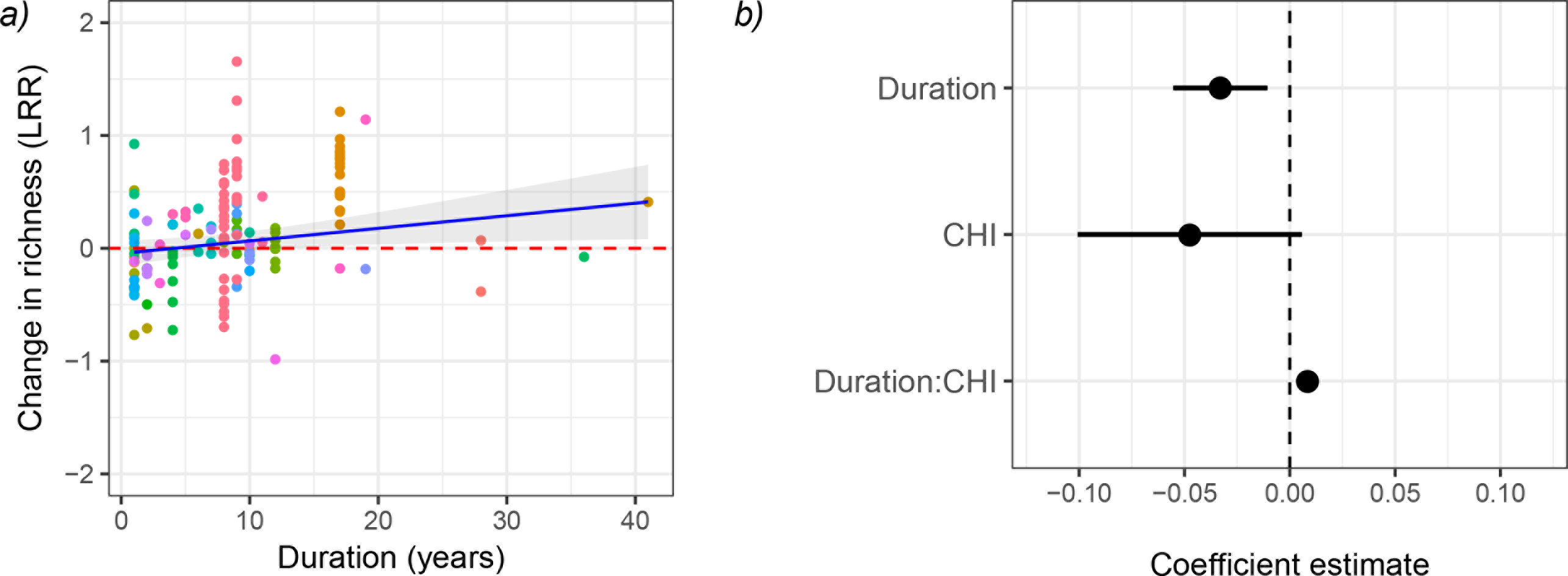
(*a*) With increasing study duration, variance-weighted meta-regression shows that the log ratio of species richness at a rate equivalent to species richness gains of 1% per year. Studies, represented by different colours, could contain data from multiple sites and so studies were modelled as random effects. (*b*) Unstandardized coefficient estimates for the relationship of the log-proportion of change in species richness as a function of study duration (Duration), shortterm cumulative human impacts (CHI; Halpern (Halpern et al. 2015)), and long-term effects of cumulative human impacts (Duration*CHI). Points represent coefficient estimates and lines represent 95% CI obtained using a variance-weighted meta-regression.′

**Table 1.** AICc scores (corrected for small sample sizes) calculated for three variance-weighted meta-regressions of the log ratio of the proportion of species richness change (LR). The model which included three specific drivers: invasion potential (Inv), nutrient addition (Nut), and rate of linear temperature change (LTC) and the model that included cumulative human impacts (CHI) both performed better than the model that did not include any form of human impacts.

| Model | Log-Likelihood | K | Deviance | AICc | ΔAICc | Akaike weight |
| --- | --- | --- | --- | --- | --- | --- |
| LR ~ Dur * (Inv + Nut + LTC) | -2492.09 | 7 | 5202.81 | 5004.06 | 0.00 | 1.00 |
| LR ~ Dur + CHI + Dur*CHI | -2497.02 | 3 | 5212.67 | 5004.64 | 0.58 | 0.85 |
| LR ~ Dur | -2518.69 | 1 | 5256.00 | 5043.61 | 39.55 | 0.00 |

Cumulative human impacts accounted for a significant proportion of the heterogeneity observed in our dataset (Q_m_ = 45.3, sites = 133, studies = 32, p < 0.001), but the directions of effect sizes were unexpected. On average, when cumulative human impacts were zero (i.e., the duration effect in Fig. 1b), the observed rate of change in species richness was −3.2% per year (95% CI = – 5.4% - (-1.1)%, p = 0.004, Appendix S1: Table S1). As expected, there was weak evidence for negative effects of cumulative human impact values on the species richness from short-term studies (i.e., the CHI effect in figure 1*b*), with an associated decline in species richness of −4.6% per unit of cumulative human impact value (95% CI = −9.6% - 0.6%, p = 0.081; table S1). However, given the interaction between impact and duration interaction (Figure 1b), each unit increase in cumulative impact value decreased the rate of species loss per year by 0.86%. (95% CI = 0.5% - 1.3%, p = < 0.001; Appendix S1: Table S1). This indicates that as cumulative human impact value increases, in long-term studies, species losses occurred more slowly than over the short-term within our dataset.

The inclusion of nutrient addition, shipping traffic (invasion potential), and the rate of linear temperature change yielded the best model (Table 1) and accounted for a significant proportion of the heterogeneity observed in our dataset (Q_m_ = 60.3, p < 0.001, sites = 110, studies = 23, Figs. 2, 3, Appendix S1: Table S1). When all specific driver values were zero (i.e., the duration effect in Fig. 2b), the observed rate of change in species richness was 1.7% per year (95% CI = 0.19% – 3.3%, p = 0.027, Appendix S1: Table S1). Importantly, different drivers had contrasting effects on local richness change when we accounted for nutrient addition, invasion potential, and rate of linear temperature change in our models. Further, when we considered the effect of these drivers on log rate of change in species richness (β_k2_), the direction of effect of each driver on local richness was reversed (Fig. 2b).

**Figure 2.**
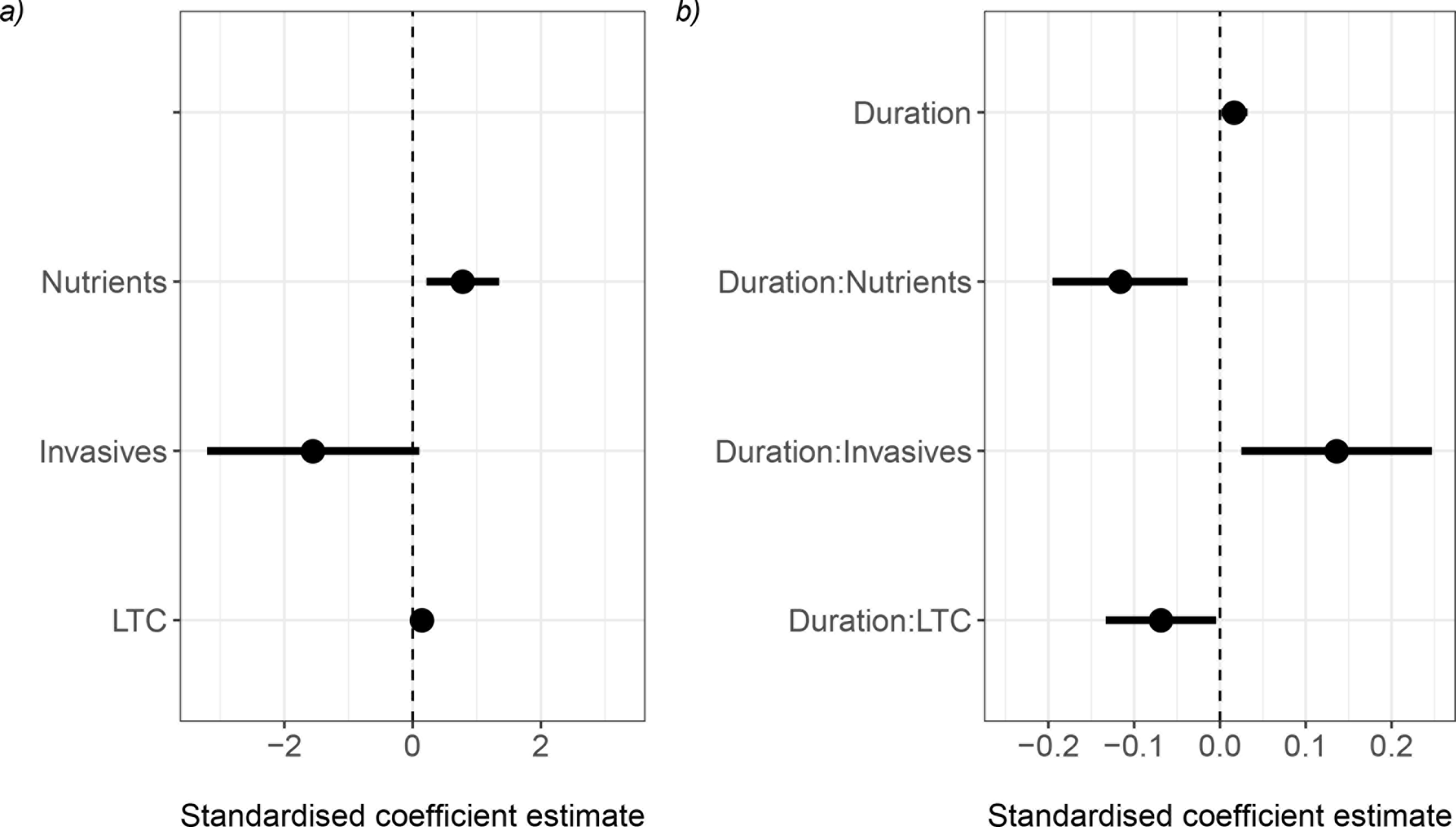
The standardized coefficient estimates of the effect of three global drivers: nutrient addition (Nutrients), invasion potential (Invasives), and the decadal rate of linear rate of temperature change (LTC) on (*a*) the log-proportion of change in species richness in the shortterm and (*b*) the effect of these drivers on the rate of change in the log-proportion of change in species richness over time. Points represent standardized coefficient estimates and lines represent 95% CI obtained using a variance-weighted meta-regression.

Nutrient addition alone was associated with increases in local gains of richness at a rate of 1.3% per tonne of nutrients / km^2^ (95% CI = 0.4% - 2.2%, p = 0.007, Appendix S1: Table S1). However, in the long-term, nutrient addition decreased the rate of species richness change over time, resulting in long-term declines in species richness (Fig. 3a). Conversely, there was weak evidence for negative effects of invasion potential (-1.4% per 1000 tonnes of shipping cargo, 95% CI = 0.4% - 2.3%, p = 0.066, Appendix S1: Table S1), but in the long-term, invasive propagule pressure positively affected the rate of species richness change over time, leading to species gains over the long-term (Fig. 3b, Appendix S1: Table S1). Meanwhile, the rate of linear temperature change was weakly associated with species gains in the short-term (6% increase per °C / decade, 95% CI = 1% – 11%, p = 0.018, Appendix S1: Table S1), but species losses in the long-term (Fig. 3c).

**Figure 3.**
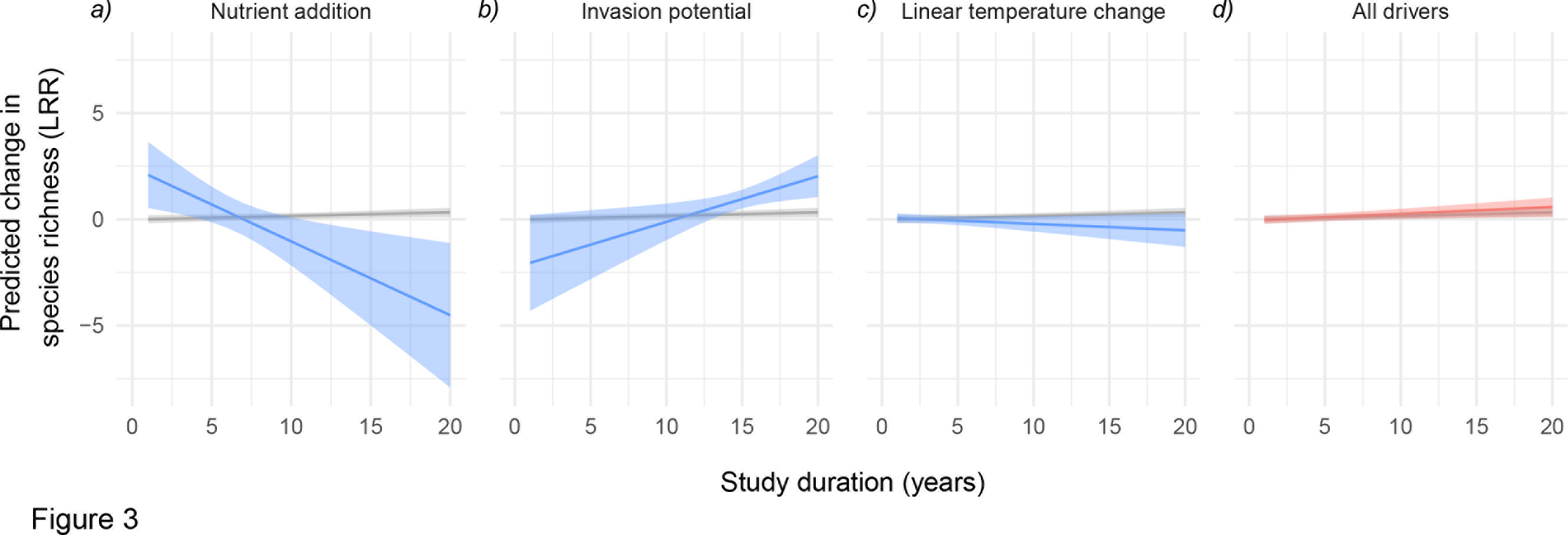
The predicted change in the log-proportion of change in species richness over study durations up to 20 years as moderated by each of the three drivers (*a*) nutrient addition, (*b*) invasion potential, (*c*) rate of linear temperature change when each is set to the maximum value observed in our dataset and the others are set to zero. The final plot (d) demonstrates the overall effect on the log ratio of local richness change when all three drivers are the maximum values observed in our dataset. Effects of drivers on predicted richness change (blue) are compared to the predicted change when all drivers are set to zero (grey). Predicted values regression lines and confidence intervals were obtained using a variance-weighted meta-regression from the full drivers model: LRR ~ Duration * (nutrient addition + invasion potential + linear rate of temperature change).

Figure 3 illustrates the expected effects of nutrients, invasive propagule pressure, and rate of linear temperature change (when set to the maximum values observed in our data) on the rate of species richness change over time compared to a baseline rate of change (1.7% per year, Fig. 3 grey line) when the three driver values are zero. The observed net effect of richness change, when all drivers were set to the maximum values observed in our dataset (Fig. 3d), shows much smaller change in the log proportion of species richness change over time compared to any individual driver. This suggests that opposing effects of local drivers can result in observations of little to no change in global averages of local richness change.

### Data coverage

With respect to global representativeness of impact levels, we had more observations of species richness change over time than expected in intermediate levels of nutrient addition and invasion potential compared to the distribution of these two drivers when considered from coastal areas globally (Appendix S1: Fig. S4a,b). Meanwhile, the cumulative human impact values ranged from 0.89 – 8.9 in our analysis, compared to minimum and maximum global values of 0 to values greater than 15. Similar to the specific drivers, the majority of our sites showed moderate impact. Fifty percent of our studies were in regions/pixels with cumulative human impact values between 2.7 and 5.1. Across taxonomic groups our data were limited to algae, fish, and invertebrate communities, or some combination of these taxonomic groups (Appendix S3). Meanwhile, temporally, eighty percent of studies were 15 years or less in duration and started after 1990.

## DISCUSSION

Our meta-analysis shows that geographic variability in human drivers can explain some of the high variability in the spatial pattern of local-scale species richness change in coastal marine systems. Furthermore, local drivers such as the nutrient inputs, invasion potential, and the rate of linear temperature change can have opposing effects on local changes in species richness. These opposing effects can interact such that the net change in local species richness can be close to zero when multiple drivers are acting on a community, as illustrated in Fig. 3d. This phenomenon may be widespread across many classes of drivers. As expected, when we considered cumulative human impacts, we observed negative effects on local richness change, though this effect was weakly supported. But, contrary to expectations, we found that cumulative human impacts were correlated with increases in the rate of change in local species richness over the long-term. This was unexpected given research showing that cumulative stressors typically have a negative effect on local communities (Christensen et al. 2006, Crain et al. 2008).

However, the cumulative human impacts are an aggregate metric; observed relationships between local scale richness change and high impacts may be driven by whatever individual driver is most important at a given location. Our results, which indicate that differences in the direction of change in local species richness can be associated with specific drivers, suggest a need to apply ecological theory about individual drivers of species richness at a local scale to broader spatial and temporal scales to generate *a priori* predictions of when and where we should observe increases or decreases in biodiversity.

### Nutrients

We found that, while sites recently associated with high nutrient run-off were associated with short-term gains in species richness, over the long-term, sites with high nutrient run-off were correlated with losses (Figs. 2, 3). Nutrient addition has been shown to increase primary production (Nielsen 2003) and richness (Bracken and Nielsen 2008) in macroalgae and may be, in part, responsible for the increase in algal richness that we observed (Appendix S3: Fig. 4). However, the processes that drive effects of nutrient addition on local communities can be complex and depend on factors such as the level of addition (Arévalo et al. 2007), species interactions (Proulx and Mazumder 1998), and dependent on time (Kraufvelin et al. 2006). Most nutrient addition studies in marine systems occur over a short time frame (e.g., Forrest and Arnott 2006, Bracken and Nielsen 2008, Herbert and Fourqurean 2008) but Kraufvelin et al. (2006) found that it could take five years before significant changes in canopy composition of rocky shore macroalgae are observed. Meanwhile in terrestrial systems, long-term studies in grasslands have also revealed that nitrogen addition can result in species losses over time (Isbell et al. 2013). Our results suggest that nutrient addition is an important driver of local richness change and that in the long-term nutrient enrichment can decrease rates of local richness change. Given the dynamics of coastal systems, this might even be more important in estuarine systems where water exchange is low relative to the open coast.

### Invasions

We found that in the short-term, there was weak evidence that recent values of shipping traffic may be associated with species richness losses, but in the long-term, high shipping traffic was associated with local gains in species richness. This observed long-term gain in species richness at sites with high shipping traffic could be the result of several processes including positive feedbacks through facilitative interactions between old and new invaders (Simberloff and Von Holle 1999, Wonham et al. 2005) or through mechanisms such as habitat change (Wonham and Carlton 2005, Thomsen et al. 2014). Our finding that long-term gains in species richness are associated with high shipping traffic is consistent with predictions made by Drake (Drake and Lodge 2004) and Sax (Sax and Gaines 2003). Elahi (Elahi et al. 2015) also found an average increase in local species richness in coastal marine communities over time, particularly for low trophic levels. When we considered separate taxonomic groups (Appendix 3: Fig. S1) we found substantial increases in the fish and algal communities. The increase in algal communities is consistent with the taxa that are transported by shipping traffic through ballast waters and organisms attached to ship hulls (Wonham and Carlton 2005).

### Temperature

Similar to the effects of nutrients on local richness, negative or low values of rate of linear temperature change over study duration, were weakly associated with short-term species gains, but high rates of temperature change were associated with long-term species losses (Figs. 2, 3c). Further this result did not appear to be strongly influenced by any one study. In a meta-analysis to examine the effects of human impacts on local species richness, Murphy and Romanuk (2013) found that increased temperature was not a significant moderator of richness in producer and ectotherm communities yet the majority of studies included in Murphy and Romanuk (2013) were less than three years in duration. Increased temperatures have been predicted to increase species richness (García Molinos et al. 2015) where warm water species move into areas that were previously at cooler temperatures at a rate that is faster than the emigration or extinction of resident species (Sagarin et al. 1999, Jackson and Sax 2010). Within our dataset, the movement of warm water fishes into areas that had previously cooler water temperatures was found in two of eleven studies examining fish communities (e.g., 53,54). However, on average, we observed that high rates of temperature change were associated with long-term declines in species richness.

### Relationship to the ongoing debate on trends in local species diversity

Broadly, our results urge caution in the interpretation of the literature on average trends in local scale biodiversity, particularly species richness, without considering local context. First, as with previous syntheses, we identified additional geographical biases in our dataset similar to those identified in terrestrial systems and in other recent syntheses of local diversity change (Vellend et al. 2013, Dornelas et al. 2014, Elahi et al. 2015, Newbold et al. 2015). Specifically, South America, Africa, Asia, and Antarctica were underrepresented. Biases of sampled sites may limit the ability to extrapolate the trends observed in our synthesis to the global scale if our dataset contains a non-representative distribution of impacts relative to all marine coastal diversity on the planet. The prevalence of drivers in our dataset differed from their global representation (Appendix S1: Fig. S4). If the same is true of other recent analyses (e.g., Vellend et al. 2013, Dornelas et al. 2014, Elahi et al. 2015), the inference of average trends in species richness could reflect spatial biases in the distribution of drivers in the datasets of these studies rather than a true global average. Recent syntheses of hundreds of space-for-time analyses report that land-use change, invasive species, nutrient addition, and habitat change are associated with declines in local-scale species richness (Murphy and Romanuk 2013, Newbold et al. 2015). When these results are translated to global maps of impacts, they suggest that richness change in terrestrial systems should be negative, on average (Newbold et al. 2015). Our results begin to attribute the magnitude and sign of local-scale species richness change to specific human impacts. Further, our results show that specific human drivers can have antagonistic effects on local richness change. We suggest a need to develop an understanding of the current and future distribution of drivers, including ones not explicitly considered in this study, to understand local species richness change across the world’s oceans. Further, we are aware of the limitation of our data in matching spatial and temporal scale of drivers with observations of species richness change and this effect of scale will need to be considered to improve the robustness of attributing human drivers to changes in local richness. We conclude that examinations of change in biodiversity that come from non-representative samples must either take drivers into account or restrict inferences to the biogeographic regions considered. This point is essential whether an analysis focuses on either temporal analysis or space-for-time substitutions.

## CONCLUSION

How global increases in species extinction rates are being manifest at local scales is relevant to basic and applied ecological research. Our analysis shows that three drivers: nutrient addition, shipping traffic, and rate of temperature change can explain some of the variability in observed trends in local marine species richness change. Our results combined with others (Murphy and Romanuk 2013, Elahi et al. 2015, Newbold et al. 2015) suggest that in coporating local context and human impacts (e.g., recent disturbances, geographic position and context of human impacts, focal taxonomic group) is more meaningful in explaining and describing patterns of biodiversity change globally than using a grand mean of local-scale diversity change. Importantly, we also demonstrate that multiple, conflicting drivers can explain some cases of no change in species richness at local scales (Vellend et al. 2013, Dornelas et al. 2014). We suggest that species richness change at local scales in coastal marine environments is an understandable and predictable phenomenon. The next step in understanding patterns of biodiversity change in the world's oceans is to incorporate the human drivers that underlie local observations of biodiversity change into syntheses of global biodiversity change. As we improve our understanding of the global distribution of human drivers, the local effects and potential interactions of these drivers we can begin to plan for the oceans of the anthropocene.

## Data accessibility

Data will be stored in the Knowledge Network for Biocomplexity (KNB) repository. Associated R scripts will be archived with the data as well as via github at https://github.com/jdunic/local-marine-metaandreferencedusingaZenodoDOI.

## Competing interests

We have no competing interests.

## Author contributions

JD and JB conceived of the study and study design. All authors contributed to data collection. JD managed the database and carried out the statistical analysis. JD and JB wrote the initial draft of the manuscript. All authors revised the manuscript.

## Acknowledgments

We would like to thank I. Myers-Smith, R. Etter, R. Stevenson, A.J. Haupt and E. Hertz for constructive comments on earlier versions of this manuscript. We thank T. Ingty for their assistance with data collection.

## Funding

We are also grateful for the financial support from MIT SeaGrant (2014-R/ RCM-36 JCD and MJSH), NatureServe (JD), Alfred P. Sloan Foundation #G-2014-13746 (JD), and the University of Massachusetts (JD, MJSH, PK). The National Science Foundation provided funding for the preparation of this manuscript (DBI-1308719 to R.E.)

